# Prediction and action in cortical pain processing

**DOI:** 10.1101/2021.09.09.459455

**Authors:** Lina Koppel, Giovanni Novembre, Robin Kämpe, Mattias Savallampi, India Morrison

## Abstract

Predicting that a stimulus is painful facilitates action to avoid harm. But how distinct are the neural processes underlying the prediction of upcoming painful events, and those taking action to avoid or prevent them? In this fMRI experiment, we addressed this by investigating brain activity as a function of current and predicted painful or nonpainful thermal stimulation, and the ability of voluntary action to affect the duration of the upcoming stimulation. Participants (*n* = 30) performed a task which involved the administration of a painful or nonpainful stimulus (S1), which predicted an immediately subsequent very painful or nonpainful stimulus (S2). On *action-effective* trials, pressing a response button within a specified time window during S1 reduced the duration of the upcoming stimulation in S2. On *action-ineffective* trials, pressing the button had no effect on upcoming stimulation. Predicted pain increased activation in regions including anterior cingulate cortex (ACC), midcingulate cortex (MCC), and insula; however, activation in ACC and MCC depended on whether a meaningful action was performed, with MCC activation showing a direct relationship with motor output. Region-of-interest analyses revealed that insula’s responses for predicted pain were also modulated by potential action consequences, especially in the left hemisphere, albeit without a direct relationship with motor output. Taken together, these findings suggest that cortical pain processing is not specifically tied to the sensory stimulus, but instead depends on the consequences of that stimulus for sensorimotor control of behavior.

**Significance statement:** During acute pain, the processing of an acute sensory event likely occurs in parallel with predictive processing about its relevance for current and upcoming voluntary behavior. Here, we temporally separated the functional processes underlying current and predicted pain and found that activation in regions typically implicated in acute pain is modulated both by the noxious nature of upcoming events and by the possibility to affect those events via voluntary action (a button press). Our findings suggest that cortical pain processing is not specifically tied to the sensory stimulus, but instead is processed in “consequence-level” terms based on what the stimulus implies for sensorimotor control of behavior.

## Introduction

The cerebral cortex plays a major role in generating predictions about internal and external events, and comparing and adjusting those predictions against incoming sensory information (Clark, 2013). But prediction is of limited usefulness if it does not ultimately lead to changes in behavior that can support the well-being and survival of the organism. The ability to predict a potentially damaging painful event is vital when that pain—and, thus, any ensuing harm—is avoidable through voluntary action. In this study, we investigated how the cortical processing of acute pain is influenced by whether it predicts a second painful stimulation in the near future, and whether participants can curtail this upcoming pain via voluntary action.

A growing body of evidence has shown that cortical pain processing and subjective experience are shaped by a variety of cognitive, affective, and contextual factors. A key observation has been that pain-related brain regions are active *before* a stimulation occurs (Atlas & Wager, 2012; Ploghaus et al., 1999; Tu et al., 2020), and can even predict whether or not it is perceived as painful; for example, prestimulus activity in anterior insula (AI) is greater for stimuli rated as painful than those rated as nonpainful (Ploner et al., 2010; Wiech et al., 2010). Expecting high (vs. low) pain also increases subjective pain reports, and this effect is mediated by activity in anterior cingulate cortex (ACC), anterior insula (AI), and thalamus (Atlas et al., 2010). This and other evidence indicate that the brain’s predictions about the potential consequences of an acute painful event influence the brain’s processing of those events.

Like expectation or prediction, action also has an integral functional link with pain. An essential feature of pain is that it motivates action to avoid harm (Morrison et al., 2013). In keeping with this, acute pain speeds reaction times (Perini et al., 2013) and facilitates specific muscle responses (Neige et al., 2018). In other words, pain is in large part an action problem rather than a sensation *per se*. In this perspective, the brain flexibly adapts its responses to current and anticipated circumstances, taking into account the available options for meaningful behavior, with cortical pain processing modulated according to relevant aspects of the behavioral context. These aspects include voluntary action selection (Perini et al., 2013) and the behavioral relevance of the stimulus as it relates to the current situation (Perini et al., 2020).

Among such flexible cortical processes, the mid-anterior cingulate cortex (MCC) plays a key role in supporting voluntary action to avoid harm. It is well-situated for this role through its involvement in the cortical control of voluntary movement during nonpainful (Amiez & Petrides, 2014; Hoffstaedter et al., 2013; Matelli et al., 1991; Procyk et al., 2016; Vogt & Morecraft, 2009; Vogt & Sikes, 2009) and painful (Misra & Coombes, 2014; Pereira et al., 2010; Perini et al., 2013, 2020; Shackman et al., 2011; Vogt, 2005) stimulation in humans and nonhuman primates. In humans, MCC’s response to pain hinges on whether an action (e.g., a button press) is performed. In contrast, AI responses to pain occur regardless of overt action (Perini et al., 2013), but have nevertheless shown a sensitivity to the behavioral relevance of the pain (Perini et al., 2020). Moreover, these two regions work together to generate subjective motivational feelings during pain, with functional connectivity between AI and MCC increasing with higher self-reported urge to move during pain (Perini et al., 2020).

If prediction and meaningful, motivated action are critical component functions of pain, how can they be disentangled for investigation of their underlying functional neuroanatomy? To do so requires experimentally dissociating acute pain processing (what is happening now) from predictive processing (what happens next), and probing the effects of motivated action (what can be done) on these processes. We pursued this aim by using fMRI to measure hemodynamic changes during an experimental paradigm in which participants were administered acute painful or nonpainful thermal stimulation (“current”) which predicted upcoming suprathreshold painful or nonpainful stimulation (“predicted”) later in the trial.

This experimental task required participants to make a timed button-press response during the stimulation in each trial, regardless of whether the current stimulus was painful or nonpainful. At the start of each trial, participants were informed of the current stimulus’ (S1) predictive relationship to an upcoming stimulus (S2), as well as whether a successfully-timed button-press within a 450ms window would shorten the upcoming stimulation by 3 seconds (“effective action” or “ineffective action” trials). This paradigm allowed dissociation of immediate from predicted effects of pain processing (current and predicted pain during S1), as well as any general effects of action from task-related processing (button-presses which would or would not affect S2). It also allowed dissociation of the effects of action execution (button-press) from action consequence (shortening upcoming stimulation).

With this paradigm, we tested for any pain-specific neural modulation of, and any interactions among, three major factors during the “current” stimulation in S1: 1) current pain in S1, regardless of whether it predicts upcoming pain in S2; 2) predicted pain in S2, regardless of current stimulation in S1; and 3) the ability to affect S2 by making an accurately-timed action during S1. In (1), *predicted* pain is held constant, potentially revealing activations selective for current stimulation; a lack of modulation here might indicate that current stimulation has no specific influence on predictive pain processing. Likewise, in (2), *current* pain is held constant, potentially revealing activations selectively modulated by the predicted stimulation. A lack of modulation here might indicate that current pain processing outweighs predicted pain.

Experimental separation of these factors pulls apart in time functions that may normally occur simultaneously during acute pain. In this perspective, a “prediction” about pain is not necessarily a prediction about the future, but can also be an in-the-moment anticipation of the potential consequences of the pain. This would serve a more general function of facilitating behavioral responses that are more likely to aid escape or avoidance, and more likely to be adapted to the particular circumstances of the painful event.

We hypothesized that AI would distinguish between predicted painful and nonpainful stimulation, regardless of whether or not the upcoming stimulus was controllable by action, whereas MCC would selectively respond to predicted stimulation that was controllable by action, regardless of whether or not it was painful. The methods, analysis plan, and specific hypotheses for each contrast of interest were preregistered on the Open Science Framework (https://osf.io/8p7rq).

## Materials and methods

### Participants

Forty participants were recruited from a student subject pool at Linköping University, Sweden using the Online Recruitment System for Experiments in Economics (ORSEE; Greiner, 2015). Inclusion criteria were the following: age range 18–40 years old, right-handed, no magnetic metals in body, no preexisting neurological history (e.g., injury, stroke), not be taking medication related to neurological or psychiatric disorder (e.g., epilepsy, depression), and absence of claustrophobia. Participants gave written informed consent in accordance with the Declaration of Helsinki and were compensated at an hourly rate of 200 SEK (approx. 20 USD). The study was approved by the regional ethics committee (Dnr 2014/340-31). Three participants withdrew from the study before any fMRI data had been collected due to discomfort in the scanner environment and seven participants were excluded due to technical problems, leaving 30 participants (17 male, age = 24.33 ± 3.25 years [*M* ± *SD*]) in the final sample. Due to technical issues, two participants only have data from one of two functional runs.

### Pain stimuli

Physical pain was delivered using a 3 × 3 cm thermal stimulator probe (Pathway model ATS, Medoc Ltd., Ramat Yishai, Israel) on the dorsal part of the left forearm. Prior to the experiment, thresholds for warmth, cold, and heat pain as well as heat pain limits were determined using a procedure adapted from Perini et al. (2013), in which the thermode had a baseline temperature of 32°C and increased or decreased at a speed of 1°C/s until participants pressed a mouse button positioned in their right hand when they felt a difference in temperature (warmth and cold thresholds; 4 trials each), when they started to feel pain (heat pain threshold; 4 trials), and when the temperature reached their maximum tolerable temperature (pain limit; 4 trials). The temperature never exceeded 50°C. After they had pressed the button, the temperature returned to baseline. The resulting pain thresholds and pain limits were used in the experiment for *painful* and *very painful* stimuli, respectively, with a maximum temperature of 49°C and at least 2°C difference between painful and very painful stimuli. Thus, painful stimuli ranged from 42°C to 47°C (*M* = 44.86, *SD* = 1.77) and very painful stimuli ranged from 45°C to 49°C (*M* = 47.86, *SD* = 1.41). Nonpainful stimuli corresponded to baseline temperature, 32°C.

### Experimental design

The experiment had a 2 × 2 × 2 within-subjects design with the following factors: S1 (current stimulation: painful or nonpainful), S2 (predicted stimulation: very painful or nonpainful), and action (effective or ineffective). The trial structure is shown in Figure 1. Each trial involved the administration of a painful or nonpainful stimulus (S1), followed by a very painful or nonpainful stimulus (S2). An experimenter (M.S.) was positioned beside the scanner and followed sound cues delivered via headphones (not audible to the participant) indicating the timing of thermode onset and offset for manual stimulus delivery. The thermode was applied to the dorsal part of the left forearm and was preprogrammed to reach the target temperature before it was applied and to remain at target temperature until it was removed. Participants’ task was to press a response button (4-Button Diamond Fiber Optic Response Pad, Current Designs Inc., Philadelphia, USA) using their right index finger as fast as they could upon presentation of a response cue (a dot), which appeared 1s following onset of S1 (1s before offset). The cue remained onscreen for 450 ms (determined based on a pilot study, see Supplementary Materials). On *action-effective* trials (50% of all trials), pressing the response button within the 450 ms time window reduced the duration of S2 from 4s to 1s. On *action-ineffective* trials, pressing the response button had no effect on the duration of S2. Participants were instructed to press the response button on all trials, regardless of whether doing so would have an effect on the upcoming stimulus. At the start of each trial, participants were presented with a brief (3s) instruction screen (see example in Figure 1), informing them about the predictive relationship between S1 and S2 (*painful* or *nonpainful*) and whether pressing the response button would influence the upcoming stimulus (*response button: enabled* or *disabled*) on that trial.

**Figure 1.**
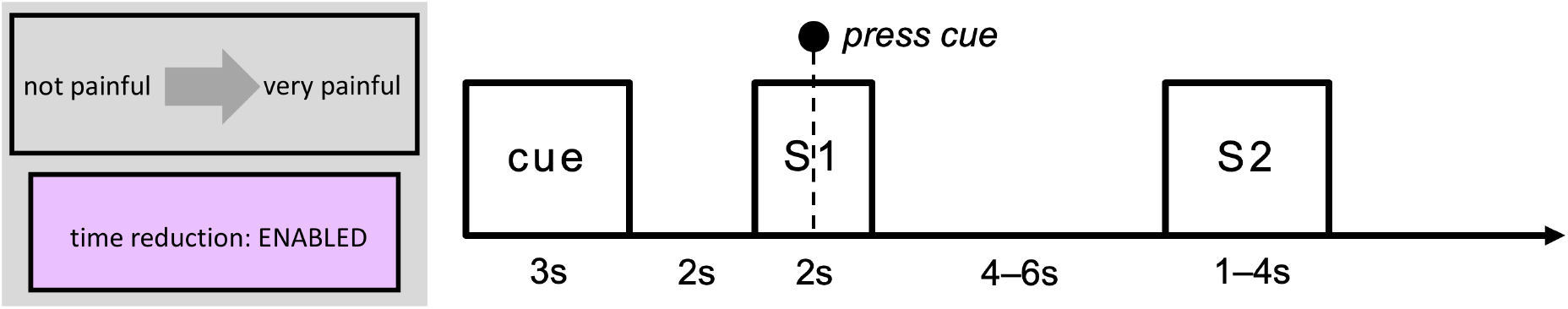
Trial structure. Each trial began with an instruction screen (“cue”; 3s), informing participants about the predictive relationship between S1 and S2 and whether pressing the response button would affect S2. A fixation cross (2s) then appeared onscreen, followed by the delivery of a painful or nonpainful stimulus (S1; 2s). Participants’ task was to press a response button upon the presentation of a cue (a dot), which was displayed 1s following the onset of S1 for a duration of 450 ms. S1 was followed by a jittered ISI of 4–6s with a fixation cross display, then by suprathreshold painful or nonpainful stimulation in S2 (1s or 4s, depending on action effectiveness and participants’ response). ITI duration varied depending on the duration of the trial and the ramping time of the stimulating thermode (ITI range 3.56–16.86s). S1 = stimulation 1 (“current pain”), S2 = stimulation 2 (“predicted pain”), ISI = interstimulus interval, ITI = intertrial interval.

The experiment was programmed in Matlab and included 96 trials, separated into two blocks of 48 trials each. The order of the two blocks was counterbalanced between participants. The order of trials within each block was pseudorandomized using a Latin square (and its reverse) balancing the four combinations of S1 and S2, to help avoid first-order carryover effects. The action factor was folded in as an additional Latin square (and its reverse). This resulted in “mini-blocks” in which the task changed every four trials. The start of each trial was triggered by the Pathway. The interstimulus interval (ISI; i.e., time from S1 offset to S2 onset) was 4–6s, randomly jittered by the Matlab script. The intertrial interval (ITI; i.e., time between S2 offset and Cue onset) varied depending on trial duration and Pathway ramping time (ITI range 3.56–16.86s).

### fMRI data acquisition

fMRI data were acquired using a 3.0 Tesla Siemens scanner (Prisma; Siemens, Munich, Germany) with a 64-channel head coil.^1^ We collected data in two functional runs (one for each task block), each lasting 20 minutes and 51 seconds. For each run, 1389 T2*-weighted echo-planar images (EPIs)^2^ containing 45 multiband slices were acquired (repetition time [TR]: 901 ms; echo time [TE]: 30 ms; slice thickness: 3 mm; matrix size: 64 × 64; field of view: 476 × 476 mm^2^; in-plane voxel resolution: 3 mm^2^; flip angle: Ernst angle [59°]). Three dummy volumes were acquired before each run (automatically determined by the system) to ensure that data collection started after the longitudinal magnetization reached steady state. A high-resolution 3D T1-weighted (MP-RAGE) anatomical image was acquired before the first EPI (TR: 2300 ms; TE: 2.36 ms; flip angle: 8°; field of view: 288 × 288 mm; voxel resolution: 0.87 × 0.87 × 0.90 mm; plane: sagittal; number of slices: 208).

### fMRI data preprocessing and analysis

MRI preprocessing and statistical analysis was performed with the Analysis of Functional Neuro Images (AFNI) software v20.0.12. BOLD images were de-spiked and slice-time corrected. For motion correction and co-registration purposes each EPI volume was registered to the volume with the minimum outlier fraction (using the AFNI outlier definition). Functional images were then warped to MNI template space using a combination of affine and non-linear transformations. Nuisance effects due to head motion (estimated from the motion correction procedure) were accounted for by adding the motion parameters as regressors of no interest in the main regression. A motion censoring threshold of 0.3 mm per TR was implemented in combination with an outlier fraction threshold of 0.05.^3^ Volumes violating either of these thresholds were subsequently ignored in the time-series regression.

A general linear model (GLM) analysis was performed to capture differences across conditions. Whole-brain, voxel-wise GLM statistical analysis was carried out on the BOLD time-series data using 3dmvm in AFNI. We conducted GLM-based analysis of a 1s time window from the onset of S1. Predictors (convolved with a standard model of the hemodynamic response function) were created for each of the eight combinations of current stimulation, predicted stimulation, and action effectiveness. Following Perini et al. (2013), we only included trials on which participants responded within 200–800 ms; all other trials were labelled as missed response and were included as a regressor of no interest. Regressors of no interest were also created for the rest of the duration of S1 (all eight conditions combined into one regressor), Cue (one regressor for each of the eight conditions), the interstimulus interval^4^ (ISI; one regressor for predicted pain and one regressor for predicted nonpain), and S2 (one regressor each for the first second of pain, the first second of nonpain, and the rest of the duration of S2 [pain and nonpain combined; this was only relevant if action was ineffective or if the participant responded slower than 450ms, otherwise the duration of S2 was 1s]). Again, these regressors only included trials on which participants responded within 200–800 ms; all other trials were captured by an additional regressor of no interest (one regressor each for S1, Cue, ISI, and S2).^5^

The AFNI program 3dClustSim was used to determine cluster-size thresholds necessary for identifying effects significant at α = .05 family-wise-error corrected. Average spatial smoothness estimates, across all participants, used by 3dClustSim were obtained using the 3dFWHMx function with the ACF flag, as per current recommendations from the maintainers of AFNI. For each analysis, we report brain areas that were activated at a voxel-wise *p*-value threshold of *p* = .002. Because AFNI outputs a single peak coordinate for each surviving cluster, a custom script was used to extract the coordinates for the first 10 peaks with the highest *t*-scores for each cluster.

In addition to the whole-brain analysis, we performed a region-of-interest (ROI) analysis in MCC and left and right insula. For the insula, we used activation clusters from a main effect of pain during S2 (see Table 3 and section “Exploratory analyses” below), excluding any voxels not in left or right insula for each respective ROI (remaining ROI cluster sizes: 257 voxels in left insula and 221 voxels in right insula). For the MCC, because there was no activation in this region in the analysis of activation during S2, we created a sphere with 10 mm radius based on coordinates reported in Perini et al. (2013). Specifically, we used coordinates from a conjunction analysis indicating common activity between motor responses during painful and nonpainful stimulation (MNI Neurosynth coordinates: –1, 8, 40, transformed from Talairach coordinates reported in Perini et al., 2013). Any voxels not in grey matter were removed (remaining ROI cluster size: 146 voxels). For each ROI, we extracted the ß values and performed a repeated-measures 2 × 2 × 2 factorial ANOVA. We also performed correlational analyses between response times and mean ß values in the MCC and left and right insula. Finally, we extracted ß values in each of the three ROIs above, as well as in ACC, during the presentation of the Cue at the start of each trial (3s), during the first second of S1, and during the first second of S2. For the ACC ROI, we used an activation cluster from the main effect of upcoming stimulation during S1 (see Table 1) and removed any voxels not in ACC (remaining ROI cluster size: 84 voxels).

**Table 1.**
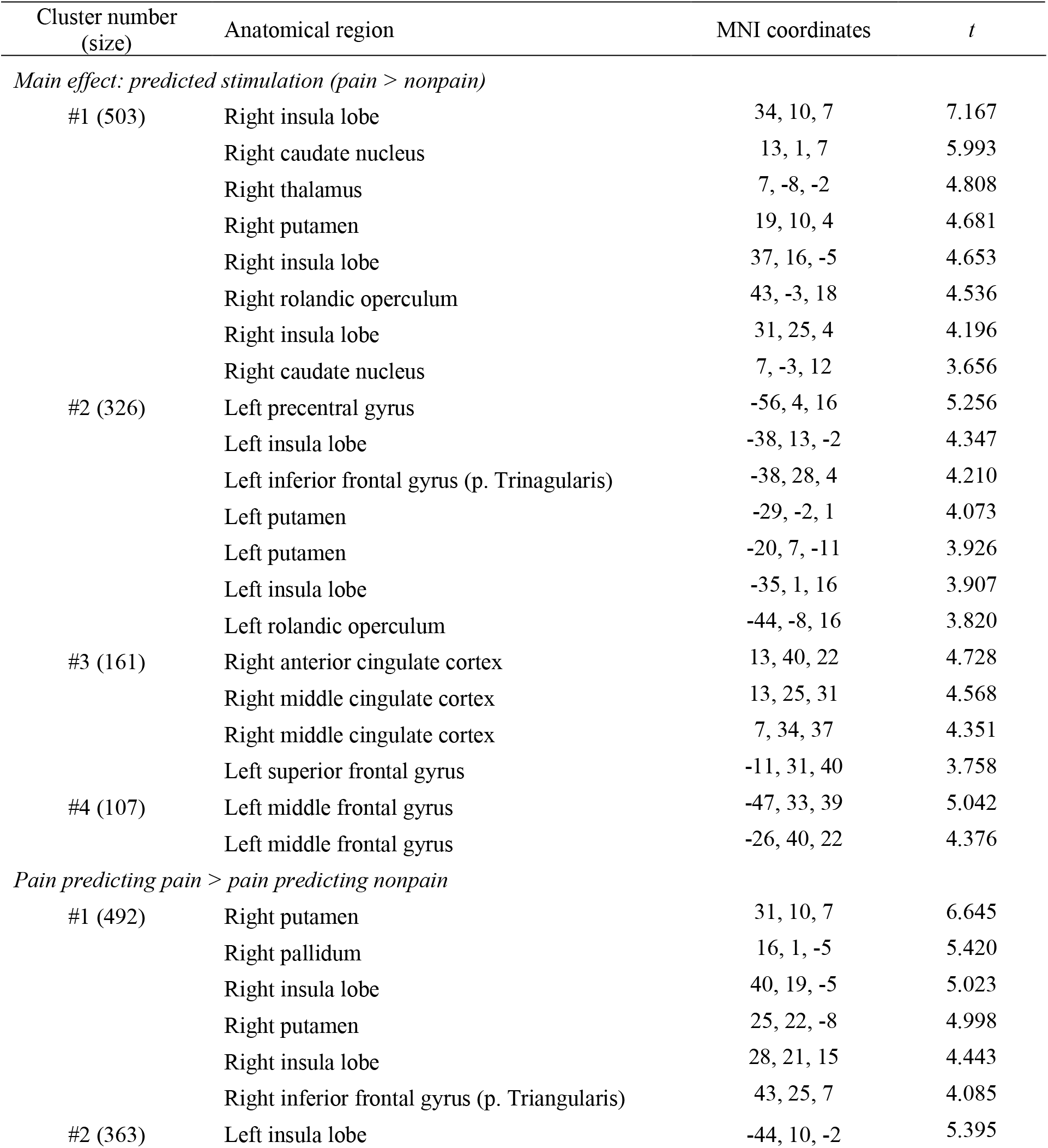
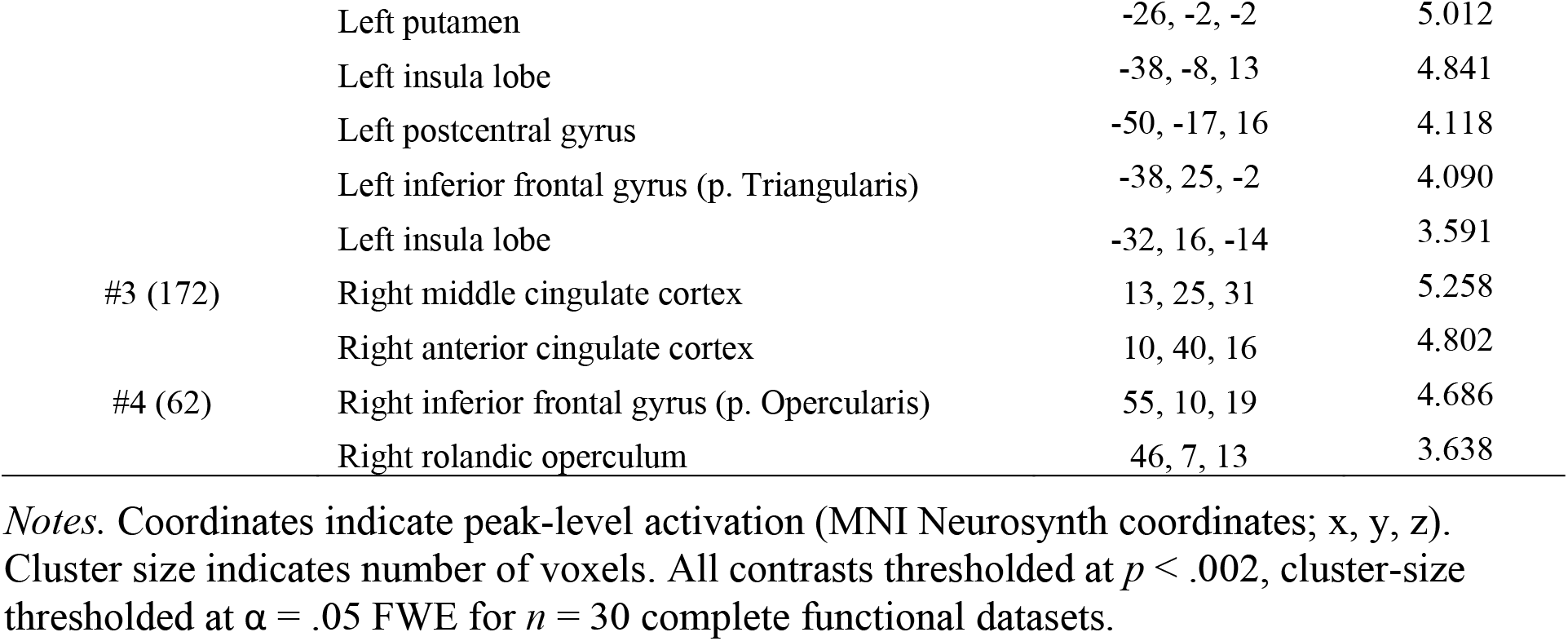
Activations from ANOVA and t-tests.

## Results

### Behavioral findings

On average, participants responded within the 450 ms response window (and thus successfully reduced the duration of the upcoming stimulation, when the response button was enabled) on 85% of trials. To investigate whether there were any task-related effects on response times, we performed a 2 × 2 × 2 repeated-measures ANOVA with current stimulation (pain or nonpain), predicted stimulation (pain or nonpain), and action (effective or ineffective) as within-subjects factors and response time as dependent variable. Only responses within 200–800 ms were included. This analysis revealed a significant main effect of action, *F*(1,29) = 19.15, *p* < .001, η_p_^2^ = 0.398, indicating that participants responded faster when the button-press action was effective (*M* = 322 ms, *SE* = 11) than when it was ineffective (*M* = 340 ms, *SE* = 10; see Figure 2). There were no main effects of current or predicted stimulation and no significant interactions, all *p*s > .05.

**Figure 2.**
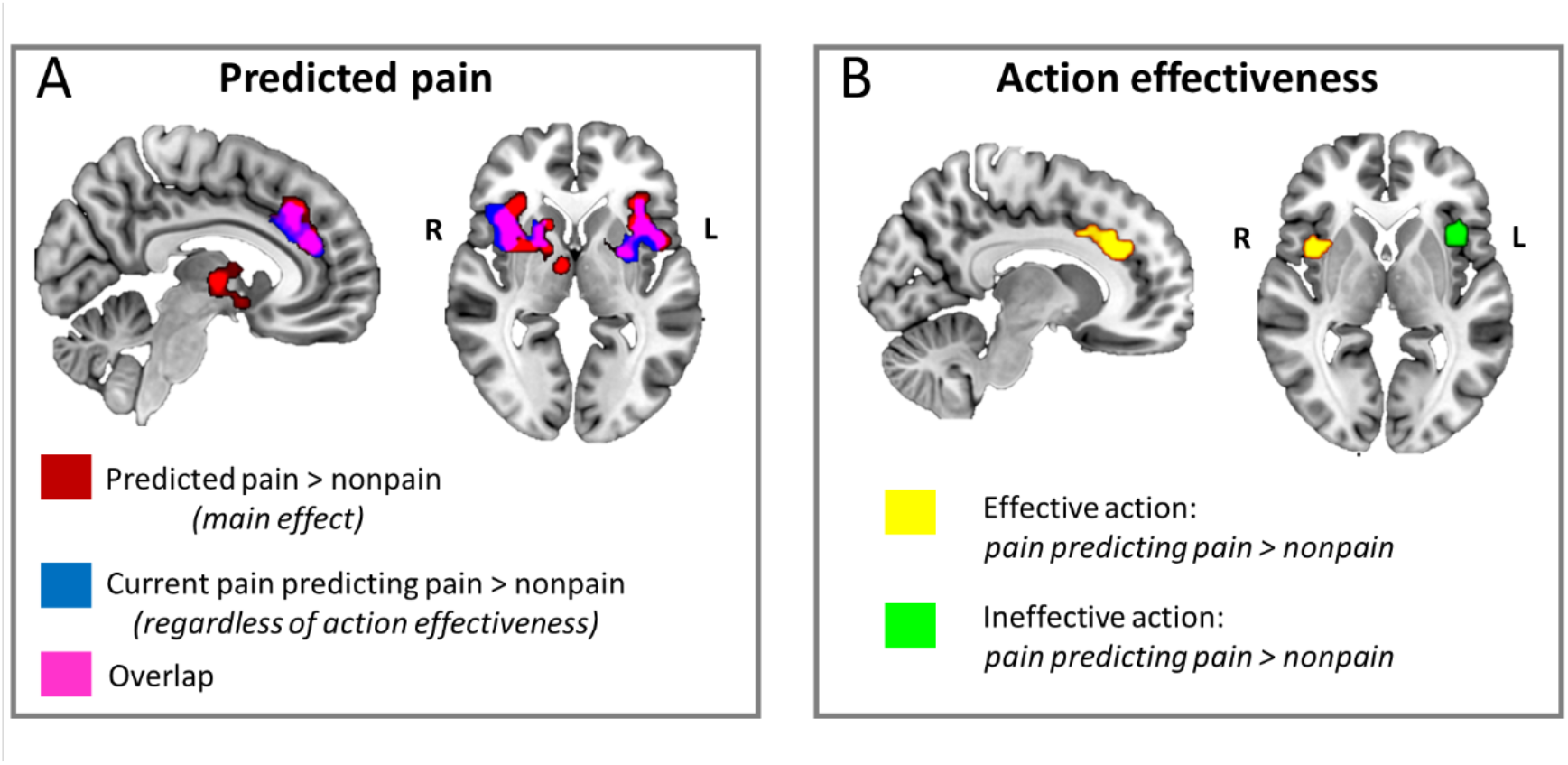
Whole-brain effects of predicted pain and action effectiveness. **A**. Noxious and innocuous thermal stimulation during S1 (“current pain”) which predicted upcoming pain in S2 (“predicted pain”) gave rise to selective BOLD increases in ACC/MCC, bilateral anterior/mid-insula, putamen, and thalamus. Red indicates clusters showing increased signal change for the main effect of predicted pain vs predicted nonpain, regardless of current stimulation or the effectiveness of the button-press action in shortening S2 duration; blue indicates clusters showing increased signal change for predicted pain when the current stimulus was painful, regardless of action effectiveness; pink indicates overlapping voxels activated in both contrasts. **B**. Whether the button-press action during S1 was effective in shortening S2 duration modulated BOLD activation in ACC/MCC and AI. Yellow indicates clusters showing selective activations for predicted pain when the current stimulus was painful during effective action trials; green indicates activation during trials in which the button-press would have no effect on S2 duration. All contrasts thresholded at p < .002, cluster-size thresholded at α = .05 FWE for n = 30 complete functional datasets. Images are displayed in radiological convention. S1 = stimulation 1 (“current pain”), S2 = stimulation 2 (“predicted pain”), ACC = anterior cingulate cortex, MCC = midcingulate cortex, AI = anterior insula.

### fMRI results: Whole-brain analyses

We first performed a whole-brain repeated-measures 2 × 2 × 2 ANOVA using the 3dmvm program in AFNI with the following factors: current stimulation (pain or nonpain), predicted stimulation (pain or nonpain), and action (effective or ineffective). Results of any significant main effects or interactions were further explored with general linear tests.

Table 1 shows the resulting brain activations from the ANOVA. First, we expected a main effect of current pain in brain regions typically recruited during pain, including somatosensory cortices SI and SII, AI, ACC, thalamus, and prefrontal cortex (i.e., “pain matrix”). However, this analysis revealed no significant activation for the main effect of current pain during S1. Second, we predicted a main effect of predicted stimulation in AI. Indeed, upcoming pain showed greater activation than upcoming nonpain in a number of regions including bilateral insula, right caudate nucleus, right thalamus, bilateral putamen, bilateral rolandic operculum, left precentral gyrus, left inferior frontal gyrus, right ACC, right MCC, left superior frontal gyrus, and left middle frontal gyrus (see Table 1 and Figure 2); the reverse contrast revealed no significant activation. Third, we predicted a main effect of action in MCC. However, there was no main effect of action. There also were no interactions between current stimulation, predicted stimulation, and action in any brain regions.

We also performed three preregistered contrasts (*t*-tests) that were of particular interest. First, we compared activation on trials in which pain predicted pain to trials in which pain predicted nonpain, in order to discover any selective activation for pain predicted by congruent current stimulation. Here, we expected increased activation in AI. This analysis revealed greater activation for predicted pain than predicted nonpain in bilateral putamen, right pallidum, bilateral insula, bilateral inferior frontal gyrus, left postcentral gyrus, right MCC, right ACC, and right Rolandic operculum (see Table 1 and Figure 2); the reverse contrast revealed no significant activation. Second, we compared activation for trials in which nonpain predicted pain to trials in which nonpain predicted nonpain, in order to discover any selective activation for pain predicted by incongruent current stimulation. Here, we again expected increased AI activation. However, this contrast revealed no significant activation. Finally, we compared trials on which predicted stimulation was painful and action was effective to trials on which predicted stimulation was painful and action was ineffective, regardless of current stimulation. Here, we predicted increased MCC activation. However, this analysis revealed no significant activation.

### ROI results

Figure 3 shows the results from the ROI analysis. Bilateral insula ROIs were defined by activation clusters from a main effect of pain during S2, while an ROI in the MCC was created as a 10 mm sphere surrounding the peak coordinates for a conjunction analysis indicating common activity between motor responses during painful and nonpainful stimulation in Perini et al. (2013). In right insula, there was a main effect of predicted stimulation, *F*(1,29) = 25.02, *p* < .001, η_p_^2^ = 0.463, indicating greater activation when predicted stimulation was painful compared to nonpainful. There were no other main effects or interactions, although the interaction between current and predicted stimulation as well as the interaction between predicted stimulation and action were both significant at α = .10 (current × predicted stimulation: *F*(1,29) = 3.06, *p* = .091, η_p_^2^ = 0.095; predicted stimulation × action: *F*(1,29) = 2.98, *p* = .095, η_p_^2^ = 0.093).

**Figure 3.**
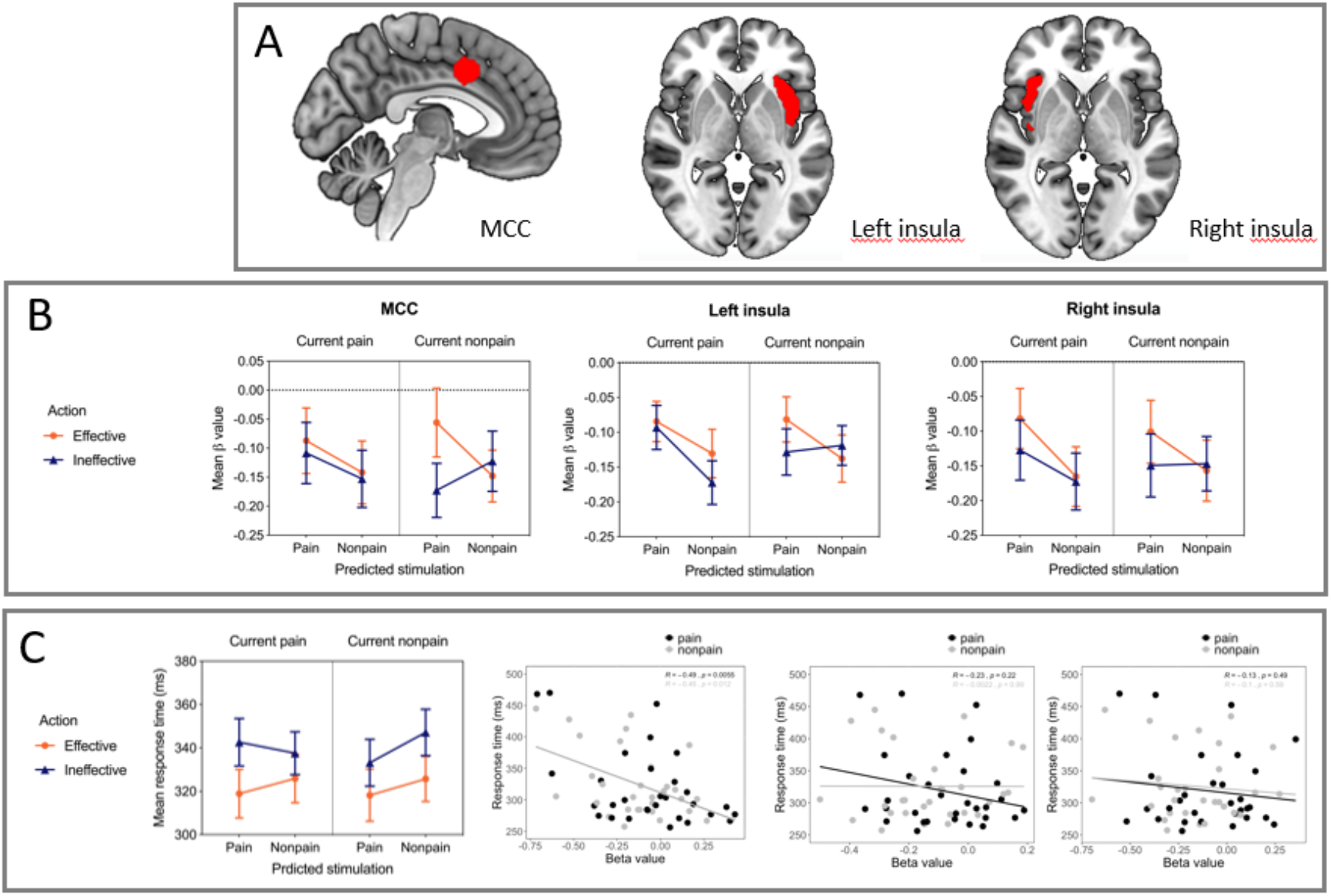
Effects of predicted pain and action effectiveness in cingulate and insula ROIs. **A**. MCC and insula ROIs: MCC cluster was defined by action selectivity during thermal stimulation in Perini *et al*, 2013; bilateral insula clusters were defined by the pain effect of painful stimulation during S2 (p < 0.002). **B**. Beta values for each ROI for the factors current pain, predicted pain, and action effectiveness. No above-threshold main effects or interactions in MCC were seen (a three-way interaction was present at an alpha level of 0.10; see text in Results section). Left insula showed a main effect of upcoming predicted stimulation (p < 0.001) and an interaction between ongoing current stimulation, upcoming predicted stimulation, and action effectiveness, (p = 0.024), with lower relative activation for trials in which predicted pain was nonpainful and action ineffective. Right insula showed a main effect of upcoming predicted stimulation (p < 0.001), indicating greater activation in cingulate and insula ROIs. **A**. MCC and insula ROIs: MCC cluster was defined by action selectivity during thermal stimulation in Perini *et al*, 2013; bilateral insula clusters were defined by the pain effect of painful stimulation during S2 (p < 0.002). **B**. Beta values for each ROI for the factors current pain, predicted pain, and action effectiveness. No above-threshold main effects or interactions in MCC were seen (a three-way interaction was present at an alpha level of 0.10; see text in Results section). Left insula showed a main effect of upcoming predicted stimulation (p < 0.001) and an interaction between ongoing current stimulation, upcoming predicted stimulation, and action effectiveness, (p = 0.024), with lower relative activation for trials in which predicted pain was nonpainful and action ineffective. Right insula showed a main effect of upcoming predicted stimulation (p < 0.001), indicating greater activation for trials in which predicted stimulation was painful. **C**. RTs and correlation with ROI beta values. An ANOVA performed on RT values with factors current pain, predicted pain, and action effectiveness revealed a main effect of action (p < 0.001), with faster RTs for trials in which the button-press action was effective in shortening S2 duration. RTs correlated negatively with MCC beta values across action trials, regardless of whether predicted stimulation was painful (r = –0.49, p = 0.005, two-tailed; see also Supplementary Figs S3 and S4), indicating a general relationship between MCC signal changes and behavioral response speed. No correlations with RTs were seen in insula ROIs (all *ps* > 0.05). Images are displayed in radiological convention. ROI = region-of-interest, RT = reaction time, S1 = stimulation 1 (“current pain”), S2 = stimulation 2 (“predicted pain”), ACC = anterior cingulate cortex, MCC = midcingulate cortex, AI = anterior insula, ANOVA = analysis of variance.

In left insula, there was a main effect of predicted stimulation, *F*(1,29) = 17.69, *p* < .001, η_p_^2^ = 0.379, and a three-way interaction between current stimulation, predicted stimulation, and action effectiveness, *F*(1,29) = 5.70, *p* = .024, η_p_^2^ = 0.164. Pairwise comparisons with Bonferroni correction indicated that activation was lower when pain predicted nonpain for which action was ineffective than when (a) pain predicted pain for which action was ineffective (*t*(106) = 3.26, *p* = .042), (b) pain predicted pain for which action was effective (*t*(103) = 3.70, *p* = .010), and (c) nonpain predicted pain for which action was effective (*t*(105) = 3.64, *p* = .012).

In MCC, there were no statistically significant main effects or interactions, although the three-way interaction between current stimulation, predicted stimulation, and action effectiveness was significant at α = .10, *F*(1,29) = 3.42, *p* = .075, η_p_^2^ = 0.105.

There was a significant negative correlation between response times and MCC activation on trials on which action was effective in shortening predicted stimulation, both when predicted stimulation was painful (*r* = –.49, *p* = .005) and when it was nonpainful (*r* = – .45, *p* = .012; see Figure 3). There was no significant correlation between response times and activation in left or right insula, neither when predicted stimulation was painful (left: *r* = –.23, *p* = .22; right: *r* = –0.13, *p* = .49) nor when it was nonpainful (left: *r* = .002, *p* = .99, right: *r* = –.10, *p* = .59). This pattern of results in MCC and insula remained regardless of whether current stimulation was painful or nonpainful (i.e., when painful and nonpainful trials were analyzed separately), although the correlation between response times and activation in the MCC when current stimulation was painful and predicted stimulation was nonpainful was only significant at α = .10 (see Supplementary Figures S3 and S4). On trials on which action was not effective in shortening upcoming stimulation, there was a significant correlation between response times and activation in MCC, but only when predicted stimulation was painful (*r* = –.39, *p* = .034; see Supplementary Figure S5); there were no other significant correlations.

### Exploratory analyses

The *t*-tests from the whole-brain analysis above revealed activation in both insula and MCC for pain predicted by congruent current stimulation (see Table 1). To further explore this result, we conducted two additional *t*-tests investigating the effect separately for action-effective and action-ineffective trials. That is, we first compared activation on trials on which pain predicted pain to trials on which pain predicted nonpain, and action would be effective upon upcoming stimulation (i.e., pain predicting “action-effective” pain vs. pain predicting “action-effective” nonpain). This analysis revealed increased activation in right insula, right ACC, and right MCC (see Table 2); the reverse contrast revealed no significant activation. We then repeated the analysis but for trials on which action was ineffective (i.e., pain predicting “action-ineffective” pain vs. pain predicting “action-ineffective” nonpain). This contrast revealed increased activation in left insula (see Table 2). We also performed corresponding *t*-tests for current nonpain. That is, we first compared activation on trials on which nonpain predicted “action-effective” pain to those in which nonpain predicted “action-effective” nonpain. This analysis revealed increased activation in right and left thalamus and right caudate (see Table 2). We then compared activation on action-ineffective trials in which nonpain predicted pain to action-ineffective trials in which nonpain predicted nonpain. This analysis revealed no significant activation.

**Table 2.** Exploratory t-tests of activation during S1.

| Cluster number<br>(size) | Anatomical region | MNI coordinates | *t* |
| --- | --- | --- | --- |
| *Effective action: pain predicting pain > pain predicting nonpain* |  |  |  |
| #1 (67) | Right insula lobe | 40, 7, 7 | 4.531 |
| #2 (61) | Right anterior cingulate cortex | 10, 31, 28 | 5.771 |
|  | Right middle cingulate cortex | 10, 13, 34 | 3.945 |
| *Ineffective action: pain predicting pain > pain predicting nonpain* |  |  |  |
| #1 (73) | Left insula lobe | -41, 16, 1 | 4.669 |
| *Effective action: nonpain predicting pain > nonpain predicting nonpain* |  |  |  |
| #1 (102) | Right thalamus | 10, -11, 16 | 5.469 |
|  | Left thalamus | -2, -8, 4 | 4.553 |
| #2 (55) | Right caudate | 13, 16, 4 | 5.263 |

*Notes.* Coordinates indicate peak-level activation (MNI Neurosynth coordinates; x, y, z). Cluster size indicates number of voxels. All contrasts thresholded at $p < .002$ , cluster-size thresholded at $\alpha = .05$ FWE for $n = 30$ complete functional datasets.

We performed an exploratory ANOVA of activity during a 1s time interval from the onset of S2, to investigate BOLD changes during delivery of an expected stimulus (S2), preceding the feedback on the outcome of the button press (shortening the stimulation). The resulting activation is shown in Table 3. There was a main effect of S2, indicating greater activation for pain compared to nonpain in bilateral insula, bilateral putamen, right pallidum, left thalamus, and midbrain; the reverse contrast revealed greater activation in left paracentral lobule, right postcentral gyrus, and left precuneus. There was also a main effect of action, indicating greater activation when action was effective than when it was ineffective in a number of regions, including insula but excluding cingulate (see Table 3). The reverse contrast revealed no significant activation. Finally, there was a significant interaction between S1, S2, and action in right superior medial gyrus, left ACC, left superior medial gyrus, right fusiform gyrus, right lingual gyrus, right calcarine gyrus, left mid orbital gyrus, bilateral middle temporal gyrus, right medial temporal pole, right inferior temporal gyrus, and left inferior occipital gyrus.

**Table 3.** Exploratory ANOVA of activation during S2.

| Cluster number<br>(size) | Anatomical region | MNI coordinates | *F* |
| --- | --- | --- | --- |
| *Main effect of pain in S2 (pain > nonpain)* |  |  |  |
| #1 (575) | Right insula lobe | 31, 13, 13 | 36.377 |
|  | Right putamen | 25, 19, −8 | 35.145 |
|  | Right pallidum | 16, 7, −5 | 23.514 |
|  | Right pallidum | 16, −5, −5 | 21.752 |
|  | Right insula lobe | 40, −5, 7 | 20.041 |
|  | Right insula lobe | 37, −17, 10 | 17.039 |
| #2 (442) | Left insula lobe | −35, 10, 7 | 41.462 |
|  | Left insula lobe | −41, 1, 7 | 36.029 |
|  | Left putamen | −20, 10, 1 | 16.785 |
| #3 (148) | Left thalamus | −8, −23, −5 | 41.128 |
|  | Midbrain | 4, −29, −17 | 29.306 |
| *Main effect of nonpain in S2 (nonpain > pain)* |  |  |  |
| #1 (83) | Left paracentral lobule | −5, −32, 70 | 22.231 |
| #2 (73) | Right postcentral gyrus | 49, −8, 31 | 19.831 |
|  | Right postcentral gyrus | 64, −5, 25 | 19.411 |
| #3 (68) | Left precuneus | −8, −56, 19 | 18.393 |
|  | Left precuneus | −14, −53, 31 | 16.714 |
|  | Left precuneus | −8, −56, 43 | 13.925 |
| #4 (61) | Right postcentral gyrus | 52, −20, 61 | 35.883 |
| *Main effect of action in S2 (effective > ineffective)* |  |  |  |
| #1 (2605) | Right supramarginal gyrus | 58, −23, 22 | 88.722 |
|  | Right rolandic operculum | 55, 13, 1 | 69.186 |
|  | Right insula lobe | 34, −17, 7 | 62.138 |
|  | Right insula lobe | 31, 28, −2 | 58.941 |
|  | Right rolandic operculum | 46, 1, 10 | 50.999 |
|  | Right middle frontal gyrus | 37, 43, 7 | 46.613 |
|  | Right insula lobe | 40, −2, −2 | 40.837 |
|  | Right inferior frontal gyrus (p. Triangularis) | 49, 25, 10 | 39.865 |
|  | Right superior temporal gyrus | 61, −47, 13 | 39.719 |
|  | Right supramarginal gyrus | 67, −41, 31 | 37.233 |
| #2 (1473) | Left supramarginal gyrus | −65, −23, 28 | 62.672 |
|  | Left rolandic operculum | −59, 4, 7 | 38.273 |
|  | Left insula lobe | −32, 13, 10 | 34.896 |
|  | Left insula lobe | −38, 1, 13 | 30.369 |
|  | Left insula lobe | −44, −2, −2 | 28.076 |
|  | Left insula lobe | −32, 28, 7 | 27.001 |
|  | Left middle temporal gyrus | −56, −62, 7 | 24.637 |
|  | Left insula lobe | –38, 7, –5 | 24.480 |
|  | Left superior temporal gyrus | –62, –50, 16 | 21.271 |
|  | Left inferior parietal lobule | –59, –35, 55 | 19.731 |
| #3 (470) | Right SMA | 7, 22, 52 | 54.180 |
|  | Right SMA | 4, 7, 52 | 32.766 |
|  | Right superior medial gyrus | 10, 40, 46 | 30.052 |
| #4 (414) | Left cerebellum (VIII) | –32, –56, 53 | 32.579 |
|  | Left cerebellum (Crus 1) | –20, –74, –29 | 18.702 |
|  | Left cerebellum (VIII) | –17, –77, –44 | 16.300 |
|  | Left cerebellum (VI) | –32, –53, –29 | 16.146 |
|  | Left cerebellum (VIII) | –20, –71, –53 | 15.843 |
|  | Left cerebellum (Crus 1) | –47, –59, –32 | 13.809 |
|  | Left fusiform gyrus | –38, –50, –17 | 13.479 |
|  | Left cerebellum (VI) | –29, –41, –32 | 13.148 |
| #5 (136) | Right postcentral gyrus | 31, –32, 73 | 36.210 |
|  | Right postcentral gyrus | 22, –44, 70 | 33.548 |
| #6 (97) | Left inferior frontal gyrus (p. Triangularis) | –38, 40, 13 | 33.992 |
|  | Left middle frontal gyrus | –47, 43, 19 | 27.496 |
| #7 (65) | Right superior frontal gyrus | 16, –2, 76 | 37.688 |
| *Three-way interaction in S2 (S1 × S2 × action)* |  |  |  |
| #1 (286) | Right superior medial gyrus | 4, 61, 13 | 26.548 |
|  | Right superior medial gyrus | 7, 58, 34 | 21.298 |
|  | Left anterior cingulate cortex | –8, 31, 13 | 17.487 |
|  | Left anterior cingulate cortex | –11, 43, 13 | 17.316 |
|  | Left superior medial gyrus | –11, 61, 28 | 15.476 |
|  | Left superior medial gyrus | –14, 55, 4 | 14.782 |
| #2 (264) | Right fusiform gyrus | 37, –65, –14 | 23.628 |
|  | Right lingual gyrus | 16, –68, –8 | 21.726 |
|  | Right lingual gyrus | 19, –77, –14 | 21.072 |
|  | Right calcarine gyrus | 19, –80, 4 | 14.153 |
| #3 (149) | Left mid orbital gyrus | –8, 40, –11 | 27.273 |
| #4 (102) | Left middle temporal gyrus | –62, –11, –14 | 21.215 |
|  | Left middle temporal gyrus | –62, –29, –5 | 14.020 |
| #5 (77) | Right medial temporal pole | 40, 7, –35 | 17.395 |
|  | Right middle temporal gyrus | 52, 7, –26 | 16.801 |
|  | Right temporal pole | 43, 16, –17 | 16.539 |
|  | Right inferior temporal gyrus | 34, 7, –47 | 14.856 |
| #6 (74) | Right inferior occipital gyrus | –29, –83, –11 | 24.242 |
|  | Left inferior occipital gyrus | –20, –92, –8 | 13.703 |

*Notes.* Coordinates indicate peak-level activation (MNI Neurosynth coordinates; x, y, z). Cluster size indicates number of voxels. All contrasts thresholded at $p < .002$ , cluster-size thresholded at $\alpha = .05$ FWE for $n = 30$ complete functional datasets.

We also visualized the BOLD signal change across conditions and trial events (cue, S1, and S2) within the ACC and insula ROIs defined by the main effect of pain described above, alongside the MCC ROI defined by Perini et al. (2013; see Figure 4). This probed any condition-selectivity across the whole trial. This showed nonselective above-baseline activation for the prestimulus cue followed by nonselective below-baseline activation for S1 in all ROIs. Selective activation emerged in S2, with bilateral insula showing a preference for painful stimulation and cingulate regions showing a preference for action trials, with highest responses for painful stimulation during action trials (post-action) in each region.

**Figure 4.**
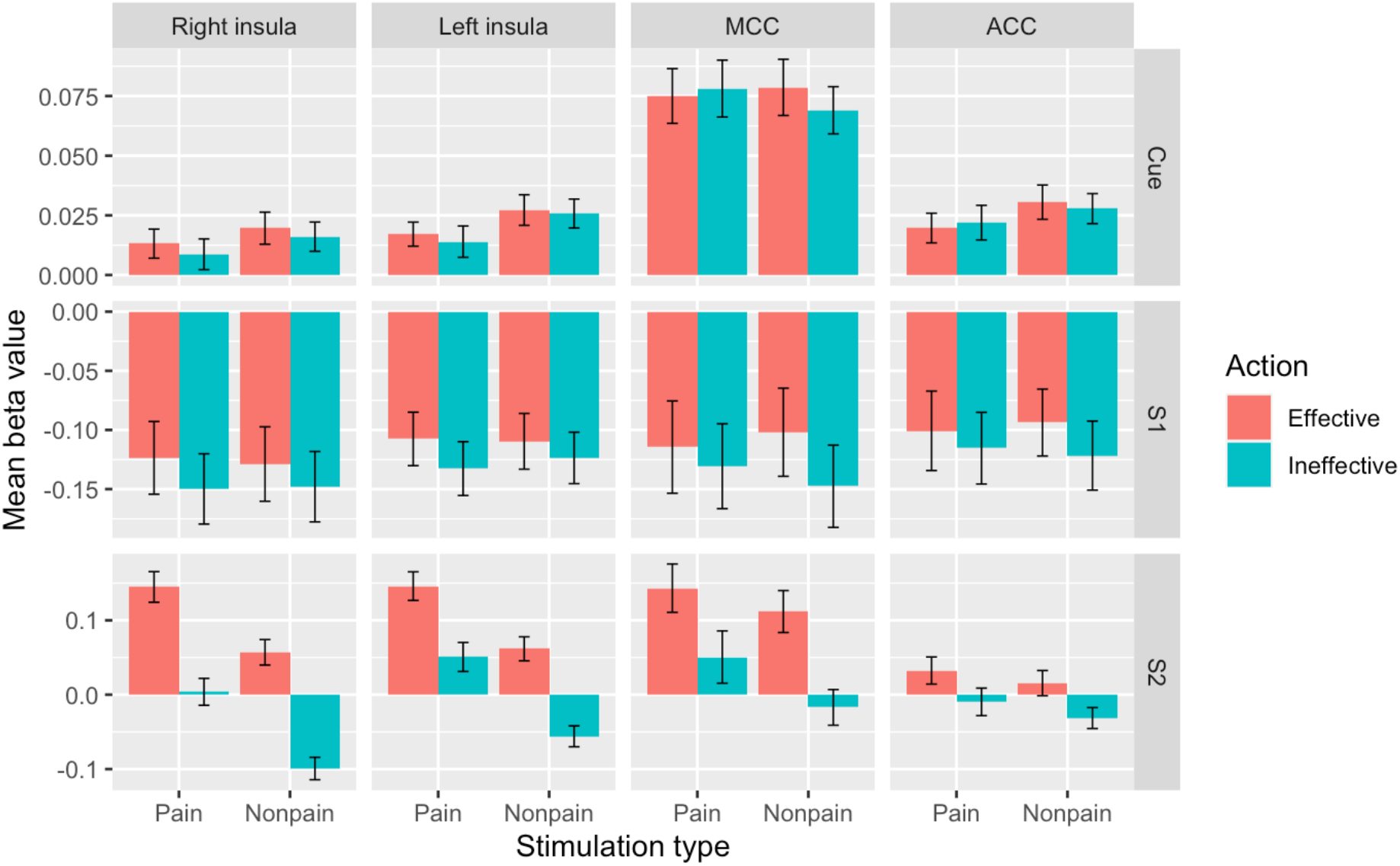
Visualization of mean beta values for thermal stimulation in cingulate and insula ROIs across cue, S1, and S2 trial components. *Top row:* cue indicating relationship between S1 and S2 elicited nonspecific above-threshold activation across conditions in all ROIs. *Middle row*: S1 elicited below-threshold activation across conditions in all ROIs (for prediction-selective responses see text and Fig 3). *Bottom row:* S2 elicited preferential activation for painful stimulation in bilateral insula, and preferential activation for action effectiveness in cingulate ROIs. ROI = region-of-interest, S1 = stimulation 1 (“current pain”), S2 = stimulation 2 (“predicted pain”).

Finally, we examined the group-level activations on unsmoothed data for the planned ANOVA and t-tests described above, as well as the exploratory t-tests. Spatial smoothing has several benefits, including improved signal-to-noise ratio, but it also has several drawbacks, such as reduced spatial resolution and potential attenuation of small meaningful activations. Reporting the unsmoothed data could therefore provide additional information, particularly regarding the spatial organization of predictive and action-related responses in the cingulate. The rostralmost activation was most selective for predicted pain regardless of either current stimulation or action, with a more caudal adjacent cluster selective only for predicted pain during current pain regardless of action. The caudalmost cluster was selective for predicted pain during current pain in effective action trials only, indicating a progressive rostrocaudal dependence on an action factor in the cingulate.

## Discussion

The cortex plays an important role in predicting upcoming internal and external events and in readying the body for action. Here, we investigated this prediction–action relationship during pain by studying brain activity as a function of both current and predicted painful and nonpainful stimulation, alongside the potential for action to affect upcoming stimulation. The study was designed to hold constant the effects of *predicted* pain by testing for main effects of painful vs nonpainful stimulation in S1 (regardless of whether it predicted upcoming pain in S2), allowing detection of any activations selective for current stimulation over and above the predicted stimulation. However, there was no such whole-brain main effect of pain during S1, indicating that, in this paradigm, current stimulation had little specific influence on predictive pain processing.

This study was also designed to hold constant the effects of *current* pain by testing for main effects of painful vs nonpainful stimulation in S2 (regardless of whether S1 was painful or nonpainful), allowing detection of any selective BOLD modulation by the predicted stimulation. This revealed that activation in anterior cingulate cortex (ACC), midcingulate cortex (MCC), and bilateral anterior and mid-insula was influenced by the noxious nature of predicted events over and above that of current stimulation. These regions were also modulated by the possibility of shortening the duration of upcoming stimulation in S2 by making an accurately-timed, effective action during S1. In particular, ACC and MCC activation for predicted pain depended on whether a meaningful action was performed, with MCC activation showing a direct relationship with motor output, which we had expected (https://osf.io/8p7rq). Contrary to our hypothesis, selective responses for predicted pain in anterior insula (AI) were also modulated by potential action consequences, especially in a cluster in the left hemisphere.

### Prediction and action in cingulate cortex

A cluster in ACC, extending into the rostral portion of MCC, was selectively engaged by predicted painful compared to predicted nonpainful stimulation (Fig 1; Table 1), consistent with previous reports of anticipatory and ongoing responses to pain in this region (e.g., Ploghaus et al., 1999). If the current stimulation in S1 predicted painful stimulation in S2, these regions showed greater engagement regardless of whether the current stimulation was painful or whether the button-press action would be effective in shortening S2 stimulation. This ACC/MCC cluster also showed more specific selective responses when current pain vs current nonpain predicted upcoming painful stimulation (Fig 1; Table 1). A subset of voxels in this ACC/MCC cluster was engaged by pain prediction only for trials in which a button-press action could affect future stimulation, but not for trials in which action would have no effect (Fig 1; Table 2). This suggests that pain-predictive activation in this region depended on whether voluntary action had a potential to affect the predicted outcome. However, in this paradigm no cingulate region was sufficiently action-selective to be activated above a corrected threshold for the whole-brain main effect of button-pressing in S1.

On the whole-brain level, current pain did not specifically modulate activation in cingulate cortex, implying that current stimulation had no strong, selective influence on pain processing above and beyond its meaning as a predictive cue. However, both ACC and MCC ROIs showed nonselective, above-baseline activation during the visual prestimulus cue (which displayed the predictive relationship between S1 and S2), followed by below-baseline responses across conditions in S1 (Fig 3). This suggests a greater nonspecific sensitivity to cue information relative to the ensuing sensory stimulation, relative to baseline. During S1, all three factors (current pain, predicted pain, and action) showed a trend (alpha < 0.1) for interaction in the MCC ROI, with greater modulation for innocuous vs noxious pain-predicting stimulation only in trials in which action would be effective in reducing S2 duration (Fig 2).

During S2, MCC showed generally greater activation for “effective action” conditions in which a potential effect of S1 button-press on S2 duration was likely expected by the participant (Fig 3). This pattern may shed light on the lack of a current pain main effect in the full model, in which regressors for cue, S1, and S2 accounted for variance in pain and anticipation-related responses over the whole trial. This possibility is consistent with the observation that temporal processing of reward/choice outcome trajectories in the cingulate can scale to the relevant timeline of trial events (Kolling et al., 2016). The across-trial pattern may also reflect the possibility that the relevant information lay in the prediction (cue) and its specific outcomes (S2), rather than the sensory properties of the S1 stimulus *per se*.

In macaque monkeys and humans, ACC and MCC contain premotor fields (cingulate motor areas/zones), with both output to and input from motoneurons in the spinal cord, well-situated for a role in action selection and control (Dum et al., 2009; Dum & Strick, 1996; Koski & Paus, 2000; Matelli et al., 1991; Monosov et al., 2020; Petrides & Pandya, 2006; Picard & Strick, 1996; Ruehl et al., 2021; Sewards & Sewards, 2003; Shyu et al., 2010; Vogt & Vogt, 2003). The organization of cingulate cortex, including the midcingulate zones, has been called “actotopic” (Caruana et al., 2018). Consistent with this, activation likelihood estimate (ALE) meta-analyses of human neuroimaging reports have shown overlapping activations for acute cutaneous pain, action execution, and action preparation in MCC (Perini et al., 2013), as well as for pain, negative affect, and control here and in and nearby cingulate subregions (Shackman et al., 2011).

The rostral midcingulate region has been specifically implicated in pain-motor relationships, for example tracking individual variance in motor reflex reactivity, with nearby areas tracking autonomic variance (Piché et al., 2010). Macaque ACC has also shown selective responses to noxious thermal stimuli during voluntary escape response (Iwata et al., 2005). On the perceptual level, intracranial microstimulation of human ACC results in reported feelings of urgency, rather than pain sensation (Bancaud et al., 1976; Hsieh et al., 1994). Nomenclature varies, but the predicted-pain cluster here likely corresponds to the human anterior rostral cingulate motor zone (RCZa; Amiez & Petrides, 2014; Loh et al., 2017).

Caudal to RCZa lies a posterior rostral cingulate motor zone (RCZp), corresponding approximately to the predefined MCC ROI (Perini et al, 2013). The motor- and action-related role of MCC is likely intimately linked to motivation to act through overt voluntary behavior (Morrison et al., 2013; Perini et al., 2013). For example, the subjective urge to move the hand away from a painful stimulus is disrupted in a type of congenital indifference to pain (Perini et al., 2020). Our previous work has shown that MCC activation during acute pain is related specifically to motor responses, but not specifically to pain, when motor responses during pain stimulation are controlled for (Perini et al., 2013, 2020). Pain activations in MCC also overlap with motor activations during the exertion of force by the hand (Misra & Coombes, 2014). Ongoing pain has been associated with pre-emptive changes in signaling dynamics in premotor cortex (Misra & Coombes, 2014) and in corticospinal tract excitability (Neige et al., 2018), with muscles under continual influence of descending faciliatory and inhibitory cortical interactions (Leis et al., 2000; Millan, 2002; Sambo, Forster, et al., 2012; Sambo, Liang, et al., 2012; Urban et al., 2004)

Consistent with these premotor and motor roles and with previous results (Perini et al., 2013) MCC responses in the present study correlated with RTs in a general fashion across pain and nonpain trials (Fig 1). RTs were faster during “effective action” trials, suggesting an increase in motivated responses in trials for which the button-presses would affect S2. (N.B: interactions (significant at α = .10) between action and pain had emerged in behavioral pilot studies outside the scanner, η_p_^2^ = 0.104 and η_p_^2^ = 0.175 in each pilot, respectively, but did not replicate in the main study; see Suppl. Mat.) MCC activation also correlates with RTs during mere observation of others’ pain (Morrison et al., 2006). Indeed, a range of effects of observed pain on motor, sensorimotor, and peripheral muscle responses has been demonstrated (Avenanti et al., 2006; Galang & Obhi, 2020; Morrison et al., 2007, 2012; Morrison & Downing, 2007; Valeriani et al., 2008), suggesting that the brain’s ability to predict and react to the likely sensory outcomes of pain generalizes to visual input about others’ actions and experiences. These findings indicate a readiness in the central nervous system for avoidance action, which can be not only anticipatory but also adaptively tailored to specific parameters of painful stimuli (Farina et al., 2003) and type of available behavioral response (Morrison et al., 2007; though see Galang et al., 2021) and task instruction (Galang & Obhi, 2020).

An exploratory plot of group-level, unsmoothed data pointed to a caudorostral gradient in the cingulate, going from a more action-specific caudal cluster (pain predicting pain > pain predicting nonpain in effective action but not ineffective action trials), to a more general pain-prediction cluster more rostrally (predicted pain regardless of action or current stimulus). A cluster for current pain predicting upcoming pain (regardless of action) was nested intermediately between them and extended more medially. This observation bolsters the proposal that cingulate premotor subregions work together to integrate stimulus content and current task demands to produce appropriate and timely responses (Kouneiher et al., 2009; Vogt, 2005; Wiech & Tracey, 2013). Such signaling likely involves recurrent feedback processing in a cingulate control hierarchy (Morrison et al., 2013), with less complex processing occurring more caudally (consistent with the action preference in the caudalmost cluster), and increasingly more nested, contingent, and abstract processing in the more rostral direction of the dorsal ACC (Loh et al., 2017; Morrison et al., 2013). Here, this rostralmost cluster showed highest responses for predicted pain regardless of task or current stimulation. Such rostral regions, such as the RCZa, may be enlisted when the situation involves higher levels of conditional information like those involved in the present task, such as increased task complexity (Kouneiher et al., 2009), internally generated actions (Mueller et al., 2007), decisions to shift from a default (Procyk et al., 2016), and comparison with current and predicted outcomes (Morrison et al., 2013). Rostral ACC regions are heavily interconnected with dorsomedial and dorsolateral prefrontal networks that also play key roles in in executive processing and action selection (Loh et al., 2017), in both current and prospective temporal windows (Kolling et al., 2014, 2016). In the case of pain, the cingulate may thus encode painful events not in sensory terms, but in terms of action consequences, analogously to the goal-level (Michaels et al., 2020) and hierarchical (Koechlin & Summerfield, 2007) encoding of actions in lateral premotor areas.

### Pain-related processing in AI is modulated by both prediction and action

Alongside the anterior portions of cingulate cortex, AI has been centrally implicated in acute pain processing (Duerden & Albanese, 2011; Knudsen et al., 2018). Here, bilateral AI/mid-insula was selectively engaged by pain prediction, regardless of ongoing stimulation or action, as well as when current pain predicted future pain (Fig 1). This is consistent with evidence that AI activity can predict whether a participant would classify a stimulus as painful, biasing “perceptual decisions” about pain even before the stimulus occurred (Wiech et al., 2010). Among other regions including the cingulate cortex, the AI mediates cue-related anticipatory effects on pain perception (Atlas et al., 2010). The prior expectation of high or low intensity pain can shift sensory processing and perceptual decision-making biases to the expected outcome, as reflected in speeded or slowed incorrect RTs, respectively (Wiech et al., 2014; Zaman et al., 2018). In conjunction with the periaqueductal grey (PAG) in the brainstem, AI is also modulated by trait anxiety in influencing pain reports of near-threshold stimuli (Ploner et al., 2010).

Previous findings have indicated that AI responses to pain are independent of overt action, when executing a button-press is compared to refraining from a button-press (Perini et al., 2013). Such evidence for pain-selective responses has been further refined by the observation that AI tracks the behavioral relevance of pain as a function of task, rather than responding to pain wholly independently of prevailing task requirements (Perini et al., 2020). Right AI was preferentially engaged by pain prediction for trials in which button-press action could affect future stimulation (Fig 2), alongside ACC/MCC. This whole-brain effect did not emerge for the left AI/mid-insula cluster (Fig 2). This was contrary to our original expectation that AI would be modulated by pain but not action. Further, activation in both left and right AI ROIs (defined by the main effect of pain in S2) was modulated by the action factor (Fig 3). All three factors (current stimulation, predicted stimulation, and action effectiveness) interacted in left insula, although in right insula this interaction was only statistically significant at α = .10).

Despite such influence of the behavioral relevance of S2 during S1, BOLD responses in AI did not correlate with RTs, either here or in a previous study (Perini et al., 2013). This suggests that while AI is sensitive to aspects of behavioral relevance in terms of action consequences, it is not directly related to producing the behavioral response. Nonetheless, responses to anticipated and ongoing pain in the AI and the MCC are likely interdependent. When an upcoming stimulus is threatening, the AI increases functional connectivity with MCC as a function of contextual information about the stimulus (Wiech et al., 2010). During painful stimulation, the functional connectivity between AI and MCC also covaries as a function of the subjective motivational urge to escape the painful stimulus through movement (Perini et al., 2020), implying that these regions play network-level roles in stamping pain with a subjective, motivated impetus to engage in escape behavior. The present findings further point to a role for AI in distinguishing whether a painful stimulus is relevant or irrelevant to the current task—and in this case, whether the current task has bearing on future outcomes.

More generally, the relative increases in both AI and ACC/MCC activity for effective action trials might appear contradictory to findings from a study by Salomons et al. (2004), which showed higher general activation in these regions during painful stimulation that was perceived as uncontrollable than when it was perceived as controllable. A possible interpretation of their findings is that increased AI and ACC activity reflects increased unpleasantness, and/or decreased predictability, of current uncontrollable compared to controllable pain. In the present study, the ability to change pain outcomes (in S2) via action (in S1) influenced brain activity *before* the painful experience occurred (Fig 3). Since behavioral control over pain may decrease both stimulus unpredictability and subjective distress, a prospective sensitivity of AI and ACC/MCC to these aspects of pain is not necessarily at odds with the findings of Salomons et al. (2004; see also (Brascher et al., 2016).

The proposal that controllability reduces aversiveness is supported by Limbachia et al., 2021) which used a Bayesian analysis to compare the BOLD responses of participants who operated a wheel via button-press, which either would (controllable group) or would not (uncontrollable group) reduce the threat of electric shock coupled with the movements of shapes on a visual display. Controllability was associated with decreased AI and increased PI responses, in contrast to the present study, in which both regions showed greater relative activation in the effective-action conditions. It may be relevant that BOLD changes in the AI ROIs, as for the MCC ROI discussed in the previous section, were below baseline for S1 but above baseline for cue; depending on the comparison, AI responses during S1 would appear as deactivations (Fig 3). It is also possible that the control/action variable is handled differently when a single action alternative is available to participants (effective or ineffective action on the group level) as opposed to two possible outcomes during the experimental session (effective or ineffective on a within-subject level). In addition, the control/action factor in our paradigm was included in a general linear model with regressors capturing relative signal changes preceding the effects of the action, whereas Limbachia et al.’s effects were derived from multilevel Bayesian modelling of button-presses alongside state/trait anxiety scores. Thus, the two approaches may have tapped different modulatory aspects related to the stimulation and task context, as well as interindividual covariation with self-reported anxiety.

AI has two major subdivisions, ventral and dorsal, though the connectivity profiles of these subregions are not always distinct (e.g., Kurth et al., 2010). The ventral subregion is associated with affective processing and is interconnected with key nodes of the classical limbic system such as the amygdala. The activations in the present study likely correspond to the dorsal subregion, which is associated with action and has anatomical and functional connections with parietal and cingulate networks (Kurth et al., 2010; Touroutoglou et al., 2012; Wiech et al., 2010). The larger activation clusters here also extended into putamen. Like the organization of cingulate cortex discussed in the foregoing section, the processing of nociceptive information may follow a caudo-rostral gradient in the insula (Cauda et al., 2012; Cerliani et al., 2012; Deen et al., 2011), going from more directly somatosensory-related processing in posterior insula to greater degree of integration in the AI, with gradual caudo-rostral shifts in connectivity with other cortical networks (Cerliani et al., 2012).

### Cortical processing of predicted outcomes

Most of the analysis in the foregoing discussion focused on the effects of current and predicted pain in S1, where holding current pain constant reveals any effects of predicted pain; and holding predicted pain constant reveals any effects of current pain. During S2, though, stimulation occurred after the task requirement in S1, with *predicted* pain thus becoming *current* pain. Here, the pain factor potentially reflects effects of expectation confirmation, based on the immediately preceding stimulus. Similarly, the action factor in S2 reflects a potential effect of feedback expectation, based on the action effectiveness and timing accuracy of the button-press performed seconds earlier during S1. To examine the effects of preceding (S1) pain on pain-related BOLD changes in S2, as well as any effects of the action task factor, an exploratory analysis investigated the factors “previous pain” [painful vs nonpainful stimulation in S1], “predicted pain” [painful vs nonpainful stimulation in S2], and action effectiveness post-response [expectation of an action consequence on S2 duration] (Table 3).

For the main effect of pain during S2, whole-brain regional activation was consistent with that expected from painful stimulation (Duerden & Albanese, 2011; Knudsen et al., 2018), including bilateral AI extending onto inferior frontal gyrus, as well as medial thalamus and midbrain. However, cingulate activation was not present in this contrast. This is consistent with previous results in which an action factor accounted for pain-related variance in this region (Perini et al., 2013, 2020). In this case, the S1 pain regressor included in the model might also have captured pain-related signal change, implying that there was no ACC/MCC signal increase for pain in S2 beyond that accounted for by the action factor overall, and the S2 pain factor specifically. Within the ACC/MCC cluster, however, signal changes were greater during the “effective action” trials (Fig 4), implying an influence of the action factor post-button-press.

In previous studies (Perini et al., 2013, 2020), the cingulate was sensitive to the action factor when the task was performed during the stimulation, as was the case during S1 in the present study. However, during S2 the ACC/MCC’s contribution to task performance had already been performed in the present study, and it may not have differentially tracked the action factor post-button-press. Instead, a cluster in the pre-SMA/SMA was engaged by the main effect of action at the whole-brain level. The engagement of the pre-SMA/SMA may reflect feedback monitoring (Graziano & Botvinick, 2002; Kolling et al., 2016; Mueller et al., 2007). Bilateral insula, bilateral lateral prefrontal and posterior parietal/TPJ regions, and cerebellum were also activated, which may make various contributions to the monitoring of expected task consequences.

In experimental pain paradigms in which a pain factor can be either controllable or uncontrollable, AI may be involved in both facilitatory and inhibitory signaling by virtue of its participation in a network including insular and prefrontal areas (Bromberg-Martin & Monosov, 2020; Jezzini et al., 2021). The findings of Bräscher et al. (2016) suggest that medial prefrontal (mPFC) and dorsal medial prefrontal (dPFC) regions may play a facilitating role in uncontrollable pain and an inhibitory role in controllable pain, respectively, via the AI (Brascher et al., 2016). In the present study, BOLD activation in the mPFC, including perigenual anterior cingulate (Brodmann areas 32/24), and orbitofrontal cortex (OFC) interacted among the factors pain [S1], pain [S2], and action [expectation of an effect of action on S2 duration] (Table 3). This interaction was also seen in bilateral anterior temporal cortex, left inferior frontal gyrus, and bilateral occipital cortex.

## Conclusion

This study temporally separated functional processes of sensory pain processing, stimulus relevance, and prediction in terms of action consequences. These processes likely occur in parallel during acute pain. BOLD activation in the ACC/MCC and AI/mid-insula was preferentially modulated by whether an upcoming, predicted stimulus would be painful, but not by whether a current stimulus was painful. ACC/MCC and AI/mid-insula were more strongly engaged when the upcoming stimulation could be affected by the possibility that a button-press would shorten it, but this action factor also influenced AI responses to innocuous stimulation that predicted pain. These findings imply that cortical pain processing in these and other cortical regions is not specifically tied to the sensory stimulus, but instead is processed in “consequence-level” terms based on what the stimulus implies for sensorimotor control of behavior.

## Acknowledgements

This study was supported by Distinguished Young Investigator grant FYF-2013-687 from the Swedish Research Council to I.M.

## Author contributions

IM designed the research. LK and MS performed the research. LK, GN, and RK analyzed the data. IM and LK wrote the paper.

## Data availability

Unthresholded statistical maps are available at

https://neurovault.org/collections/XBRBXVRD/

## Footnotes

1 We stated in the preregistration that a 12-channel head coil would be used; however, the 12-channel head coil was replaced with a 64-channel head coil prior to the start of data collection for our study.

2 We stated in the preregistration that 456 EPIs would be collected for each functional run; this number was based on the number reported in a previous publication using a different protocol and is an error. However, because the number of EPIs is directly linked to the duration of the experimental paradigm, this error should not affect the credibility of our findings.

3 We stated in the preregistration that fMRI data preprocessing and analysis would be performed according to current guidelines from the maintainers of AFNI. At the time, recommendations involved using a motion censoring threshold of 0.2 and an outlier fraction threshold of 0.1, as well as including the motion parameter derivatives as regressors of no interest, as mentioned in the preregistration. However, recommendations have since been updated, and changes were made to our preprocessing script based on the new recommendations. Thus, minor deviations from the preregistered preprocessing plan simply reflect updates in the AFNI guidelines.

4 The preregistration states “inter trial interval” rather than “interstimulus interval”; this is an unintentional mistake in the preregistration.

5 We did not specify in the preregistration exactly which regressors of no interest would be included in the analysis; any deviations resulting from the inclusion of regressors of no interest reflect the non-specificity of the preregistered analysis plan rather than deliberate changes.

## Notes

https://neurovault.org/collections/XBRBXVRD/

### Competing Interest Statement

The authors have declared no competing interest.

