## Supplementary materials for "Prediction and action in cortical pain processing"

### Pilot studies

In order to confirm the appropriateness of the task design, we conducted two pilot studies in which participants performed the task outside the scanner. In the original version of the task, the time from Cue offset to S1 onset was 1s and the duration of the press cue was 350 ms (with a 350ms response time window). However, because the effect on response times seemed weaker than predicted in pilot study 1, we conducted an additional pilot study in which we increased the time from Cue offset to S1 onset to 2s in order to allow more time for participants to process the instruction. Because response times were slower than anticipated, indicating a low proportion of trials on which participants succeeded to respond in time to reduce the duration of the upcoming stimulation, we also increased the duration of the press cue (and thus the time window in which participants could affect upcoming pain) to 450ms in order to increase the sense of controllability. The methods and results from each pilot study are described in detail below.

#### Pilot study 1

**Participants.** 28 participants were recruited using the same methods as in the main experiment, except they only needed to meet inclusion criteria for age (at least 18 years old) and not be taking pain relieving medication, antidepressants, anxiolytics, or other medication that may influence the perception of pain. Participants provided written informed consent in accordance with the Declaration of Helsinki and were compensated at 200 SEK/hour. One participant was excluded due to a technical error, leaving 27 participants for analysis. Due to technical issues, twelve participants only completed one of two runs of the task.

**Materials and procedure.** The pain stimuli and task were identical to those reported in the paper, with two exceptions: (1) time from Cue offset to S1 onset was 1s and (2) the duration of the press cue was 350ms. Thus, participants had to respond within 350ms in order to reduce the duration of S2 on controllable trials. Participants were seated at a desk and completed the task on a computer. The experimenter (L.K.) was positioned next to the participant and followed sound cues delivered via headphones (not audible to the participant) indicating the timing of thermode onset and offset for manual stimulus delivery.

**Results.** We performed a 2 × 2 × 2 repeated-measures ANOVA with current stimulation (pain or nonpain), predicted stimulation (pain or nonpain), and action (effctive or ineffective) as within-subjects factors and reaction time as dependent variable. Results are shown in Supplementary Figure 1. There was a significant main effect of predicted stimulation, $F(1,26) = 5.99$, $p = .021$, $\eta_p^2 = 0.187$, indicating that participants responded faster when the predicted stimulation was painful ($M = 487$ ms, $SE = 10$) than when it was nonpainful ($M = 502$ ms, $SE = 10$). There were no main effects of current stimulation ($p = .828$) or action (although the effect was in the predicted direction, $F(1,26) = 3.49$, $p = .073$, $\eta_p^2 = 0.118$) and no significant interactions (although the interaction between predicted stimulation and action was significant at $\alpha = .10$, $F(1,26) = 3.01$, $p = .094$, $\eta_p^2 = 0.104$).

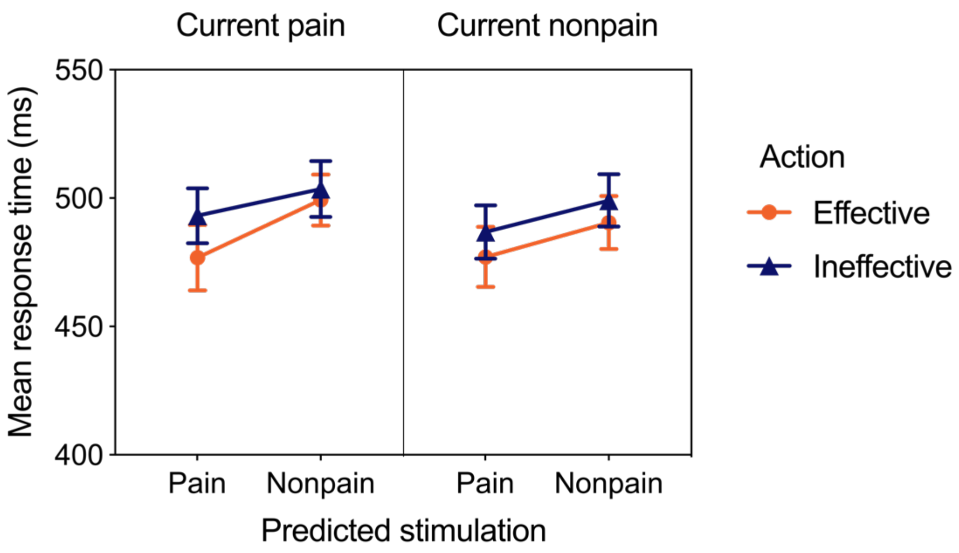


**Figure S1.** Mean response time as a function of current stimulation, predicted stimulation, and action in Pilot Study 1. Error bars represent standard errors.

### Pilot study 2

**Participants.** 21 participants were recruited and compensated as in pilot study 1. Due to technical issues, one participant only completed one of two runs of the task.

**Materials and procedure.** The pain stimuli and task were identical to those reported in the paper. Participants were seated at a desk and completed the task on a computer. The experimenter (L.K.) was positioned next to the participant and followed sound cues delivered via headphones (not audible to the participant) indicating the timing of thermode onset and offset for manual stimulus delivery.

**Results.** We performed a 2 × 2 × 2 repeated-measures ANOVA with current stimulation (pain or nonpain), predicted stimulation (pain or nonpain), and action (effective or ineffective) as within-subjects factors and reaction time as dependent variable. Results are shown in Supplementary Figure 2. There was a significant main effect of action, $F(1,20) = 7.64$, $p = .012$, $\eta_p^2 = 0.276$, indicating that participants responded faster when the button-press action was effective ($M = 492$ ms, $SE = 9$) than when it was ineffective ($M = 511$ms, $SE$

= 9). There were no main effects of current or predicted stimulation and no significant interactions, although there was a close to significant three-way interaction, $F(1,20) = 4.25$, $p = .053$, $\eta_p^2 = 0.175$. Pairwise comparisons with Bonferroni correction indicated that response times were faster when pain predicted action-effective pain than when (1) pain predicted action-ineffective pain ($p = .005$), (2) pain predicted action-ineffective nonpain ($p = .042$), (3) nonpain predicted action-ineffective pain ($p = .037$), and (4) nonpain predicted action-ineffective nonpain ($p = .026$).

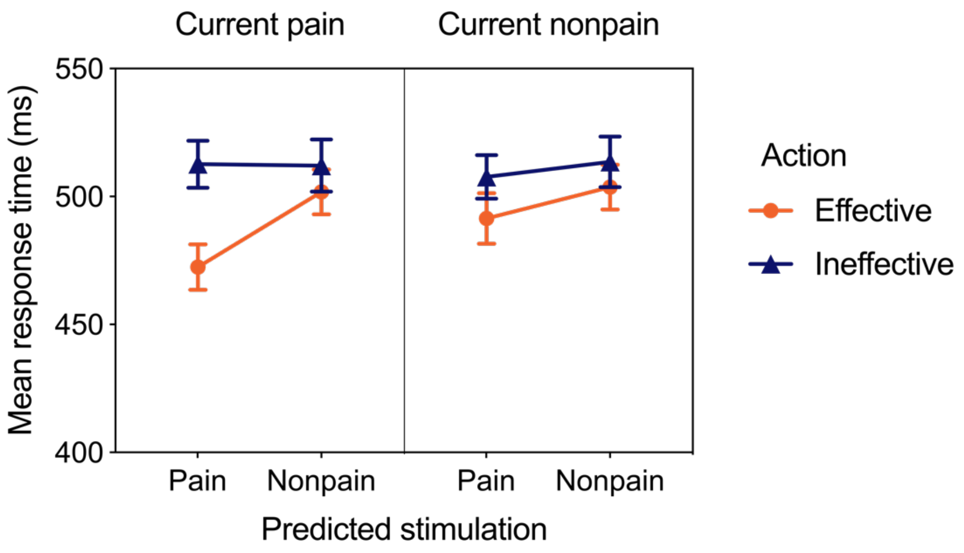


**Figure S2.** Mean response time as a function of current stimulation, predicted stimulation, and action in pilot Study 2. Error bars represent standard errors.

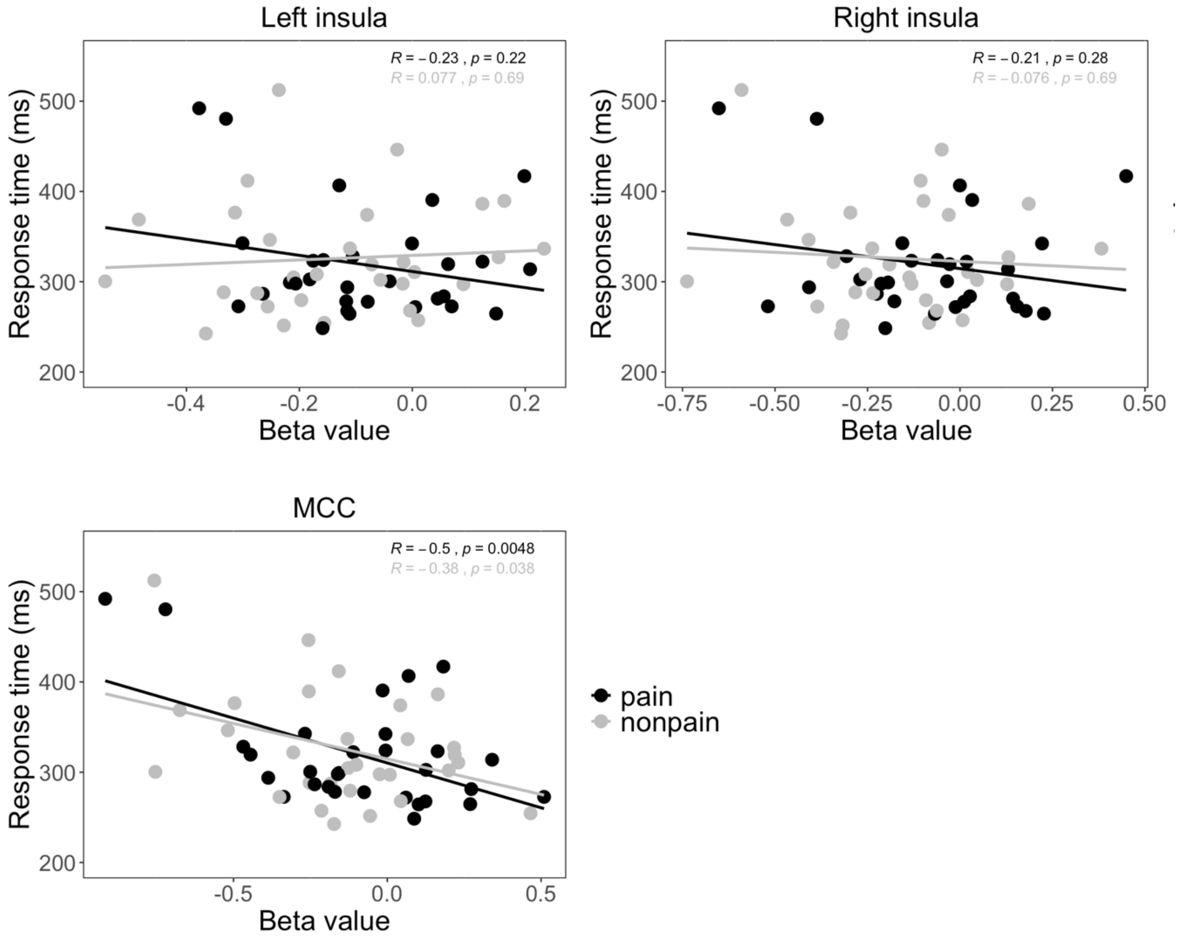


**Figure S3.** Correlations between response times for predicted painful and nonpainful stimuli and ß values of activation in MCC, left insula, and right insula on trials on which current stimulation was painful and action was effective.

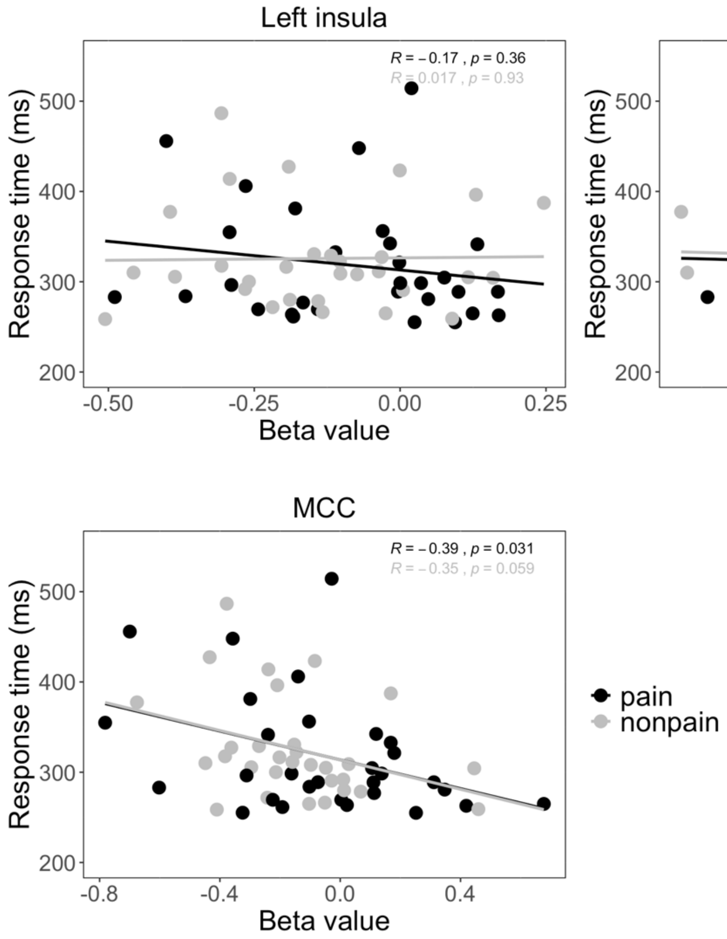


**Figure S4.** Correlations between response times for predicted painful and nonpainful stimuli and ß values of activation in MCC, left insula, and right insula on trials on which current stimulation was nonpainful and action was effective.

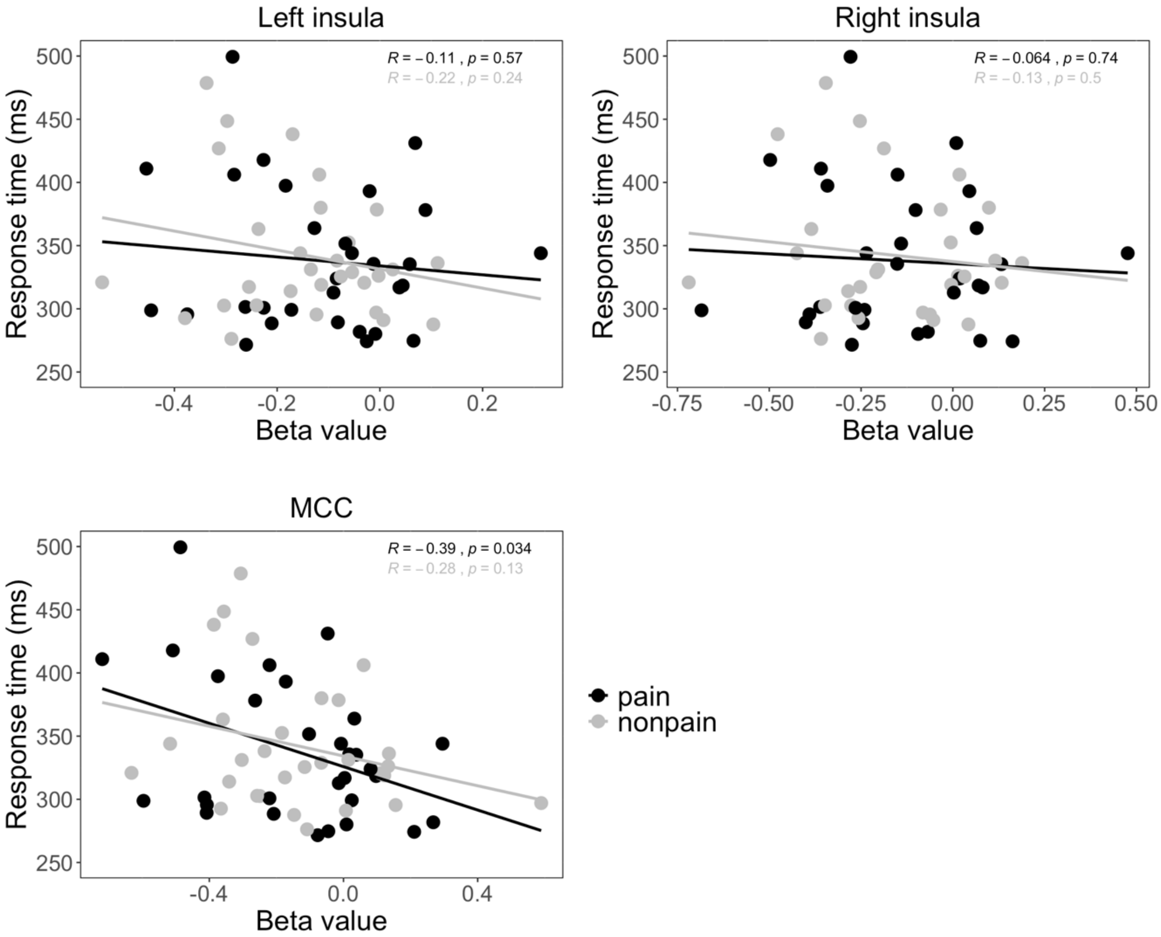


**Figure S5.** Correlations between response times for predicted painful and nonpainful stimuli and ß values of activation in MCC, left insula, and right insula on trials on which action was ineffective.
